# The Impact of Anesthesia on Ultrasonic Glymphatic Perturbation Protocol: Alertness, and Glymphatic Influx

**DOI:** 10.1101/2024.01.24.576888

**Authors:** Haijun Xiao, Binita Shrestha, Alexander Aviles Cruz, Jeffrey J Iliff, Tyrone Porter, Muna Aryal

## Abstract

We have recently discovered that transcranial low-intensity focused ultrasound can enhance glymphatic transport, which facilitates the removal of waste metabolites from the brain in a preclinical rat model [1]. The central hypothesis was that ultrasound, functioning as a pressure wave, has the potential to influence the convective forces generated by arterial pulsations, which are also pressure waves, serving as the primary mechanism for glymphatic transport. Importantly, our data revealed that (i) the ultrasound protocol (650 kHz at 0.2MPa for 10 minutes to the entire brain) is safe, as histological evaluations showed no parenchymal damage and no difference in levels of neuronal degeneration or astrocytic activation 72 hours after the intervention, and (ii) the required pressure is significantly low, 10 times below FDA-approved limits for diagnostic ultrasound, and can be achieved using existing FDA-approved clinical transcranial focused ultrasound systems, suggesting its ease of translation into clinical applications. However, the study was conducted under specific physiological states with approximately 2.5% isoflurane-induced anesthesia. There is a limitation in fully understanding the potential bioeffects of ultrasound within the glymphatic space across various physiological conditions, primarily because the glymphatic transport’s efficiency fluctuates with anesthetically induced physiological states. Hence, establishing standardized protocols of ultrasonic glymphatic transport at optimal physiological state is crucial for ensuring consistent, reliable results across research laboratories during the preclinical development of this technique. The primary objective of this study is to evaluate how ultrasonically manipulated glymphatic transport is influenced by different levels of isoflurane-induced anesthetic conditions, including 3% (just above 2.5%), 2% (slightly below 2.5%), and 1.5% (quasi-awake state, resembling the existing clinical practice of ultrasound treatment). Results show that the impact of ultrasound during glymphatic transport depends on the level of anesthesia. It increases alertness in lightly anesthetized animals and improves glymphatic transport, while it induces drowsiness in heavily anesthetized animals, leading to reduced glymphatic transport. The results suggest that lighter anesthesia is beneficial for efficient ultrasonic glymphatic transport, making it more in line with the awake state in current clinical ultrasound treatments, thus advancing the technology closer to clinical translation.

## Introduction

The brain fluid compartments are involved of intracellular fluid (60-68%), interstitial fluid (also known as extracellular fluid) (12-20%), blood (10%), and cerebrospinal fluid (10%) [2]. The blood-brain barrier and blood-cerebral spinal fluid barrier separate blood from the interstitial and cerebrospinal fluid, respectively. The cerebral fluid moves through the ventricles and subarachnoid space of the cortex and spinal cord, entering the brain parenchyma perivascularly to cleanse the brain before draining into the lymphatic system. Recent studies have found that cerebral and interstitial fluid interchange continuously, facilitated by the movement of cerebrospinal fluid along the periarterial space into the Virchow-Robin spaces, and then transported into the brain parenchyma by AQP4 water channels expressed in astrocytic endfeet. This system of convective fluid fluxes, with rapid interchange of cerebral and interstitial fluid, is called the glymphatic system and functions similarly to the lymphatic system in peripheral tissue [3]. The transport of solutes in the glymphatic system is largely dependent on the arterial pulsation [4]. Further, a dysfunctional glymphatic system can elevate the risk of developing neurovascular, neuroinflammatory, and neurodegenerative disorders. Briefly, a research study conducted a comparison of glymphatic function between young and old mice, revealing a substantial 80-90% decline in aged mice [5]. This decline in glymphatic function due to aging could contribute to the accumulation of misfolded proteins, potentially leading to neurodegenerative diseases. Investigations carried out on human subjects have established a connection between reduced glymphatic function and Alzheimer’s patients [6]. MRI measurements suggest that cerebrospinal fluid contrast agent enters the brain and is subsequently cleared, and the clearance process is delayed in patients with dementia [7]. Significant irregularities within the perivascular space extend beyond Alzheimer’s disease, encompassing non-Alzheimer’s dementia, characterized by alterations in or surrounding cerebral blood vessels due to factors such as hypertension, atherosclerosis, or hereditary conditions [8]. Consequently, the glymphatic system assumes a key role in upholding brain health and cognitive function by facilitating the elimination of waste products and toxic proteins from the brain. Different lifestyle choices are recommended for enhancing the functionality of the glymphatic pathway as protective and preventive measures against both aging and Alzheimer’s disease [9]. These choices include ensuring sufficient sleep, staying hydrated, engaging in regular exercise, maintaining a healthy diet, consuming caffeine in moderation, practicing mindfulness and stress reduction, trying intermittent fasting, creating a comfortable sleep environment, considering certain supplements, refraining from excessive alcohol and smoking, and exploring techniques like cranial manipulation and massage. The relationship between the glymphatic function and different physiological states like sleep, along with diseases such as Alzheimer’s or traumatic brain injury, is explored. However, they are based on correlations because there aren’t established methods to directly control glymphatic transport. Recent research suggests that low-level transcranial electrical stimulation can improve deep sleep (Lim et al. 2023), a process closely linked to accelerating glymphatic clearance [10]. Similarly, photobiomodulation therapy, also known as low-level light therapy, aims to enhance the glymphatic drainage system (Salehpour et al. 2022). Nevertheless, both techniques encounter a common challenge: limited tissue penetration, potentially restricting their effectiveness in targeting and influencing deeper brain structures. Therefore, there’s an urgent need to discover more effective interventions to address impaired glymphatic drainage systems.

Transcranial low-intensity focused ultrasound presents a promising tool in neuroscience for noninvasive interrogation of the central nervous system. This technique demonstrates the ability to generate bioeffects even in deeper regions of the brain, utilizing FDA-approved clinical devices. One of the applications includes temporarily loosening the blood-brain barrier, the tight junction between endothelial cells in the cerebral vasculature, enabling precise delivery of therapeutic and biological agents at a millimeter level of accuracy [11]. This effect is achieved by combining ultrasound with intravenously introduced microbubbles, typically used as contrast agents for diagnostic imaging, which help open the vascular barrier. The vascular barrier opening technique’s entry into clinical trials brings significant potential for enhancing drug delivery in the treatment of various neurological diseases and disorders [12–14]. In addition to improving drug delivery efficiency, the targeted opening of the vasculatures in a particular brain region can also influence the broader dynamics of interstitial-cerebrospinal fluid flow, a phenomenon observed in both preclinical [15,16] and clinical contexts [17,18]. Further, higher-pressure ultrasound can similarly enhance molecular movement within targeted brain regions, even without opening the vascular barriers [19]. However, it is still unclear whether utilizing lower-pressure ultrasound in a non-targeted manner even without opening the vascular barrier, could enhance the essential process of glymphatic clearance.

Indeed, we have pioneered an ultrasound method that employs low-intensity levels and can be applied comprehensively throughout the brain in a non-targeted fashion to enhance glymphatic transport effectively, all without requiring the opening of the vascular barriers. We hypothesized that ultrasound, functioning as a pressure wave, has the potential to influence the convective forces generated by arterial pulsations, which are also pressure waves, serving as the primary mechanism for glymphatic function. In brief, our technique employed 650kHz ultrasound frequency, 0.2 MPa in situ pressure, and a 7.7% local duty cycle applied throughout the brain for 10 minutes [1]. Importantly, our findings showed that (i) the ultrasound protocol is safe, as confirmed by histological evaluations showing no damage and no difference in neuronal degeneration or astrocytic activation 72 hours after the intervention, and (ii) the required pressure is notably low, being tenfold below FDA-approved limits for diagnostic ultrasound and can be achieved using existing FDA-approved clinical transcranial focused ultrasound systems, suggesting its potential for clinical use. Following our reports, which initially highlighted a specific glymphatic bioeffect within the lower-level ultrasound paradigm, others have documented similar findings [20]. However, our initial investigations, along with those conducted by Yoo et al., involved subjects under particular physiological states induced by specific anesthetics—either isoflurane (2.5%) as detailed in [1] or a combination of 40–80 mg/kg ketamine and 10 mg/kg xylazine per [20]. These conditions restrict a comprehensive understanding of the potential of ultrasonic glymphatic transport across diverse physiological contexts. It is important to know that the glymphatic transport system works differently depending on the physiological state. During sleep, the system is more effective because of increased fluid between brain cells and specific brain wave patterns, but not all types of drug-induced sleep or deep anesthesia work equally well for clearing brain waste. For example, one of the studies in whole-brain in-vivo imaging showed that the active diffusion of compounds from the cerebral spinal fluid space into the brain tissue happens during wakefulness, and the administration of general isoflurane anesthesia can negatively affect the cerebral spinal fluid circulation within the brain, especially with higher doses of the anesthetic agent [21]. Therefore, further research is required to assess the effectiveness of ultrasound under different physiological conditions to establish a standard protocol in a preclinical setting. Thus, our study aimed to investigate how varying levels of isoflurane anesthesia impact ultrasonic glymphatic transport during fluid exchange, to pinpoint the optimal physiological condition for ultrasound therapy before its application in disease and translational models. The current study was conducted under three specific isoflurane-induced anesthetic conditions: 3% (slightly above 2.5%), 2% (just below 2.5%), and 1.5% (mimicking the awake state, akin to clinical settings). Throughout the study, we administered two imaging tracers, IR800 (Free IRDye800-1kDa) and IR800-IgG (IRDye800-labeled IgG antibody-160 kDa), via intrathecal injection to get access to the glymphatic space. Two imaging tracers were used to investigate whether different anesthetic conditions played a role in the transport of tracers of varying sizes through the glymphatic pathway. This analysis was prompted by our previous research, which showed that at a 2.5% anesthetic level, the smaller tracer (IR800-1 kDa) diffused more extensively than the larger one (IR800-IgG-150kDa) [1]. The physiological states were monitored during ultrasound therapy, and the distribution of the tracers in ex-vivo brain slices was subsequently assessed.

## Materials and Methods

### Stereotactic-Guided Low-Intensity Transcranial Focused Ultrasound System

#### Calibration

Before employing the sequence for ultrasonic glymphatic manipulation, the stereotactic-guided preclinical low-intensity transcranial-focused ultrasound system was characterized in both in-vitro (Figure 1) and in-vivo settings (Supplementary Figure 1). Briefly, the ultrasound system included a 650 kHz spherically focused single-element transducer, a generator that incorporated the degassing system, a passive cavitation detector, a 3D-positioning system, and a water tank (Image Guided Therapy-in Pessac, France). The transducer was calibrated using a 1 mm needle hydrophone (Precision Acoustic in Dorchester, United Kingdom). The dimensions of the ultrasound field were assessed in terms of its full width at half maximum at the lateral and axial planes, representing the diameter and length of the ultrasound focus respectively. These dimensions measured approximately 3 mm x 20 mm, as shown in Figures 1D and E respectively. To confirm the ultrasound field’s behavior in in-vivo conditions, we implemented the vascular barrier opening assay using an established ultrasound pulse (650 kHz, 1PRF, 1 min, 0.35 MPa), in combination with an ultrasound contrast agent (Optison – GE HealthCare, Chicago, IL), as illustrated in Supplementary Figure 1. The vascular opening approach is consistent with our [22,23] and other previous papers [14,24,25], making it a widely recognized method for evaluating ultrasound fields in in-vivo settings. Furthermore, it has already been integrated into clinical practice for targeted therapy [14,26]. The vascular opening visualized the leakage of tissue dye in the targeted brain regions (identified by white circles in Supplementary Figure 1C), indicating that the lateral direction of the ultrasound field matched the diameter of the circle ∼3mm. The transducer’s axial coverage (20 mm) is extensive enough to extend across the entire brain from its anterior to posterior regions. Since the ultrasonic glymphatic manipulation protocol involves systematically exposing the entire rat brain region using ultrasound in a grid pattern, it becomes imperative to verify the precision of our sonication protocol in terms of both its targeting and coupling between the transducer and skin of the rat’s head. To achieve this, we replicated the grid pattern approach used for glymphatic application by opening the BBB. Consequently, we observed a consistent and uniform dispersion of trypan blue leakage as shown in Supplementary Figure 1E. This observation further validates that our sonication pattern for glymphatic manipulation is precisely aligned with the intended targeting and coupling parameters. It’s important to note that we utilized significantly lower pressure (0.2 MPa) in glymphatic manipulation purposes as compared to the method of BBBO (0.35 MPa) and omitted the use of microbubbles.

**Figure 1.**
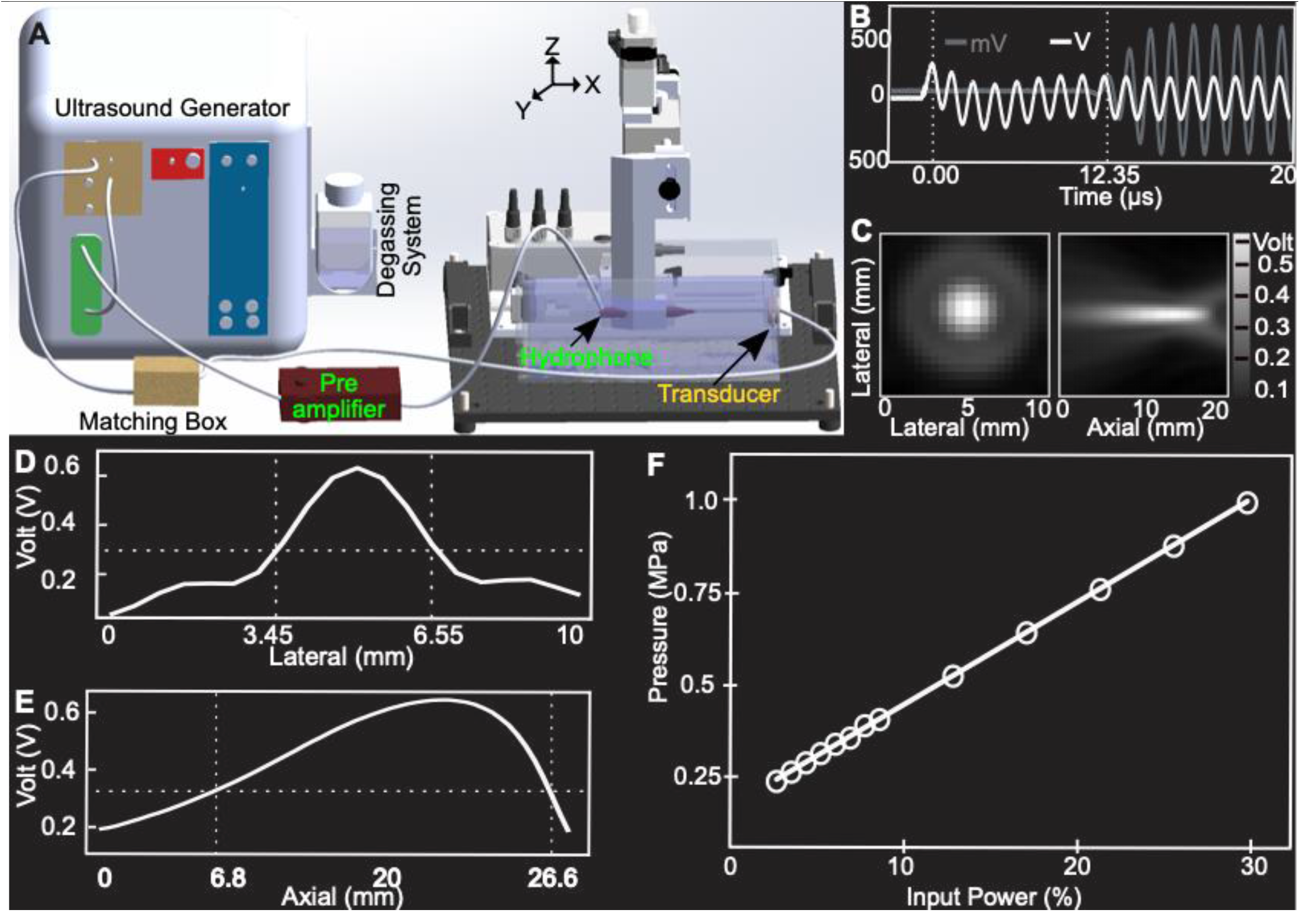
Ultrasound transducer calibration in in-vitro. **A**. Schematic of stereotactic-guided low-intensity transcranial focused ultrasound system that consists of an ultrasound generator, a 650 kHz focused transducer with matching box, hydrophone with preamplifier, and a 3D positioning system. The transducer that is submerged in a water tank receives an electric voltage from a generator. A hydrophone that is mounted on a 3D positioning system is placed within a focus of the transducer to scan the pressure field. **B**. Representative sinusoidal waves of ultrasound with ∼12.35 Μs of time of flight. **C**. Two-dimensional scan of lateral and axial planes. **D-E**. Full width at half maximum of the ultrasound field at the lateral and axial planes that represent the diameter and length of the ultrasound focus (∼3mm x 20mm). **F**. Ultrasound pressure at the focus of the transducer at different input power.

### Delivery Agents

Near Infrared region dye IRDye® 800CW NHS Ester (IR800∼1kDa) and IR800 conjugated human immunoglobulin G (IgG) antibody (IR800-IgG ∼160kDa) were used as model drugs. IR800 was purchased from LI-COR Biosciences (Lincoln, NE). The conjugation of IR800 with IgG was performed by our collaborator at the University of Texas at Austin by following the manufacturer’s protocol. Briefly, the human IgG was purchased from Sigma (Sigma Aldrich, St Louis, MO). Then the IR800 dye was dissolved in DMSO (10mg/ml). The human IgG (5mg/ml) was prepared in sodium phosphate azide-free buffer (pH 8.5). The dye was added to the IgG solution at a final ratio of 2:1. The mixture was allowed to react for two hours at room temperature in the dark. The labeled IgG was purified using a Zeba Spin® desalting column 40K (Thermo Scientific, Waltham, MA). The purified labeled IgG was collected, and shipped on dry ice to the Loyola University Chicago Laboratory, and stored at -20°C until use. The absorbance of labeled dye was measured using a UV-Vis-NIR spectrophotometer (Shimadzu, Kyoto, Japan). The degree of labeling was calculated using the following formula:

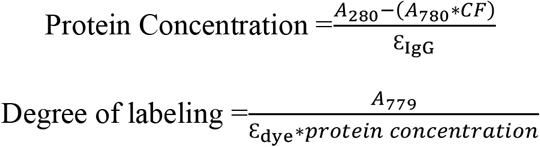

where, A_280_ and A_780_ are the absorbances of dye and IgG respectively, Ɛ_IgG_ (210,000 M^-1^ cm^-1^) and Ɛ_dye_ (240,000 M^-1^ cm^-1^) are molar extinction coefficients for the IgG and dye respectively in PBS. CF is the correction factor of 3% recommended by the manufacturer.

### Characterization of Delivery Agent (IR800-IgG)

UV-Vis-NIR spectrophotometer and Sodium dodecyl sulfate-polyacrylamide (SDS-PAGE) gel electrophoresis were used to confirm the conjugation of IR800 to IgG. For SDS-PAGE gel electrophoresis, 10 μg of Control IgG (unlabeled) and IR800-IgG (labeled) were mixed with 4X Laemmli sample buffer containing 0.05% volume β-mercaptoethanol. The mixture was heated at 90°C for 5 minutes on a heat block. The samples were loaded on 4-20% Mini-PROTEAN®TGX precast protein gels (Bio-Rad) and the electrophoresis was performed at 100 mV for 75 minutes. The gels were stained with Coomassie Blue for two hours then destained overnight. The gels were imaged using GelDoc™ EZ imager (Bio-Rad). To confirm the fluorescence signal from IR800-IgG, the gel was imaged using an Odyssey LICOR infrared imager using an 800 nm channel. Before animal use, the stock solutions of IR800 and IR800-IgG (1 mg/mL) were prepared by individually dissolving them in DMSO, which were subsequently diluted with artificial cerebral spinal fluid (CSF) to obtain a working solution (0.02 mg/mL). To compare the ultrasonic glymphatic-based diffusion, both, IR800 and IR800-IgG were administered intrathecally according to the experimental groups. The dose of each delivery agent was 4 μg/Kg, and the injected volume was 80Μl. Figure 2A shows that the IRDye ® 800CW NHS ester was conjugated to IgG through primary and secondary amines. The UV-Vis -NIR spectra of IR800-IgG demonstrate an IgG absorbance peak at 280 nm and a dye absorbance peak at 780 nm in Figure 2B. Based on these absorbance values, the degree of labeling for IR800-IgG was found to be 2.49 moles per mole of IgG. The conjugation was further confirmed using SDS-PAGE gel electrophoresis, shown in Figure 2C. The β-mercaptoethanol reduced IgG samples, both control and labeled, resulting in two bands, one from heavy chains ∼50 kDa and another from light chains ∼25 kDa. The fluorescence images of the gel show IR800 signal at both bands indicating that IR800 dyes were conjugated to both heavy and light chains.

**Figure 2.**
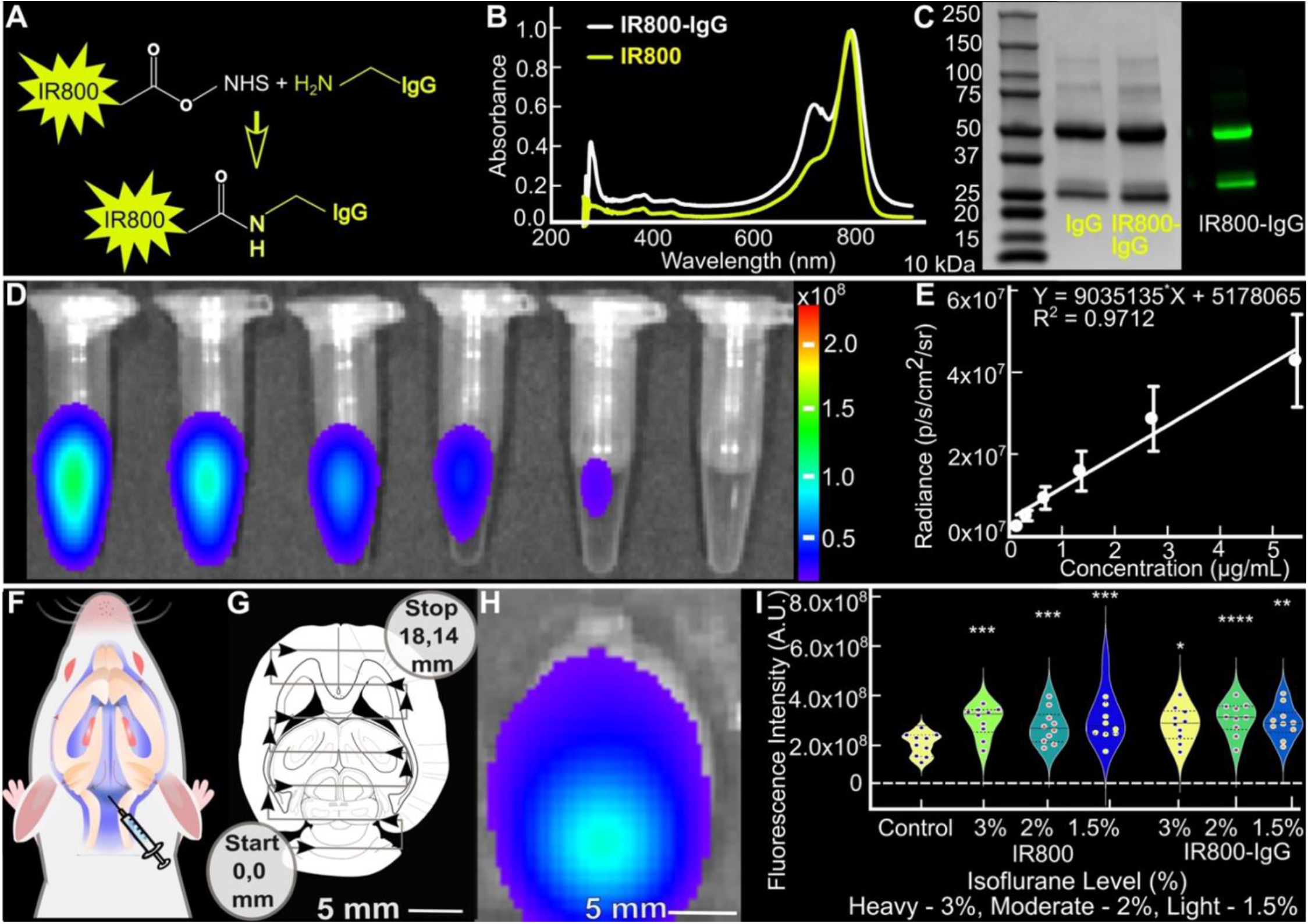
Characterization and calibration of IR800 in in-vitro and in-vivo settings for glymphatic transport analysis. **A**. Scheme illustrating IR800 dye conjugation to IgG via NHS chemistry. **B**. UV-Vis spectra of IR800 and IR800-IgG. **C**. SDS-PAGE gel images showing Coomassie blue staining (left) and fluorescence at 800nm (right). **D-E**. Calibration curve for IR800 in in-vitro settings using IVIS. **F**. Schematic depiction of intrathecal injection to access imaging tracers within the glymphatic system. **G**. Planning of ultrasound trajectory across the entire brain using rat stereotactic coordinates for glymphatic manipulation. **H**. Ex-vivo IVIS image of a rat brain demonstrating successful IR800 delivery via intrathecal injection. **I**. Calibration curve for IR800 in ex-vivo settings using IVIS, utilizing the ex-vivo image from (H) under varying anesthetic conditions. Data represented as mean ± S.D. *: p ≤ 0.05, **: p ≤ 0.01, ***: p ≤ 0.001, ****: p ≤ 0.0001.

### Animals

Healthy male Sprague Dawley rats were obtained from Charles River and used after acclimation for three days. A total of 44 rats were used in this study. Among them, 5 rats were specifically designated for in-vivo calibration involving vascular barrier opening, while the remaining 39 rats were utilized for ultrasonic glymphatic manipulation. The rats had an average weight of 273 ± 25 grams during the study. The use of animals in this research was approved by the Institutional Animal Care and Use Committee (IACUC) at Loyola University Chicago. The rats were divided into three main groups based on anesthesia levels: Heavy (anesthesia with 3% isoflurane), Moderate (anesthesia with 2% isoflurane), and Light (anesthesia with 1.5% isoflurane). Two imaging tracers of different sizes (IR800 -1kDa and IR800-labeled IgG antibody-160 kDa) were injected into the spinal fluid via cisternal magna as described below, leading to the creation of two sub-groups within each anesthesia group: a Control group and an Ultrasound group. During the experiment, a careful observation of several crucial physiological indicators in the rats was conducted, including heart rate, respiratory rate, oxygen levels, perfusion, and body temperature. This comprehensive monitoring aimed to ascertain whether any of these vital indicators might be influenced by the manipulation of ultrasonic glymphatic activity under distinct anesthetic conditions. Following a ten-minute interval from the cisternal injection, the Ultrasound group underwent a 10-minute exposure to 650kHz ultrasound at a pressure of 0.2 MPa, covering the entire brain region. In the Control group, we maintained identical experimental conditions as in the Ultrasound group, except keeping the ultrasound input at 0 MPa throughout the 10-minute sonication period. Seventy minutes after the cisternal injection, the rats were euthanized, and their brains were extracted for ex-vivo optical imaging using IVIS. This process aimed to measure the amount of the tracer delivered to the brain tissue.

### Intrathecal Cisterna Magna Injection

The animals were situated in a chamber where they received ambient air containing 4% isoflurane through an electronic vaporizer (SomnoFlo, Kent Scientific Corporation) for around 5 minutes. Once the absence of paw reflex was confirmed, the animals were weighed and secured onto a custom-made stereotaxic frame created using a 3D printer (Ender-3, Creality 3D Technology) within the laboratory. The administration of isoflurane for the animal experiments was calibrated to meet the specified target levels using a calibrated vaporizer. To maintain a consistent level of anesthesia throughout the procedure, the flow rate of isoflurane was adjusted based on the animals’ body weight. A heating pad connected to a physiological monitoring system (PhysioSuite, Kent Scientific Corporation) was positioned beneath the rats’ bodies to ensure warmth and comfort. Eye lube (OptixCare Eye Lube Plus) was applied to provide lubrication and moisture. In preparation for optimal ultrasound beam coupling and access to the cisterna magna for intrathecal injection, the hair on the rats’ dorsal scalp and 3 cm towards the neck was removed using an electric hair trimmer (Philips Norelco 5500 Series), followed by the application of Nair baby oil hair remover lotion. The specified amount of model drug solutions was then mixed with artificial cerebral spinal fluid (Tocris, Minneapolis, MN) and introduced into the cisterna magna of the rats using a 27-gauge 13 mm butterfly needle, connected to a 12 cm polyurethane tubing (SAI Infusion Technologies, Lake Villa, Illinois) and a one-milliliter syringe via a 27-gauge 13 mm needle (PrecisionGlide, BD) as noted in Figure 2F.

#### Ultrasonic glymphatic manipulation protocol

Following the cisternal injection, animals were secured within a plastic stereotactic frame and immobilized using ear bars and a bite bar, all coupled to a 3-axis positioning system with the 650-kHz transducer. Then, the isoflurane level was adjusted to 1.5%, 2%, and 3% based on the experimental groups. A thin layer of ultrasound gel was applied to enable the connection between the water-filled coupling membrane of the ultrasound transducer and the skin of the head. The sonication trajectory was determined using the transducer’s remote positioning capabilities across all three axes. The stereotactic coordinates for sonication followed a grid pattern as depicted in Figure 2G. In brief, sonication began at the START location (0 mm, 0 mm), proceeded rightward following the black arrow to reach (0 mm, 14 mm), then moved upward to arrive at (3 mm, 14 mm). Six of these 3 mm x 14 mm sub-loops constituted the trajectory leading to the STOP point of the sonication (18 mm, 14 mm), effectively covering the brain of an adult rat. The entire duration for completing one trajectory loop was approximately 27.12 seconds, with a 50 ms pause time between each loop, resulting in a total of 30 loops for each rat. Throughout the process, the ultrasound remained continuously active while the transducer gradually swept the ultrasound trajectory, taking approximately 14 minutes for the completion of 30 loops. For sham procedures, the identical positioning and trajectory were employed, but the input power to the ultrasound transducer was zero. So, ultrasound, at an in-situ pressure of 0.2 MPa with a duty cycle of around 7.7%, was applied transcranially across the entirety of the brain for 14 minutes. Considering skull attenuation, a 30% pressure insertion loss was assumed for rats of this size and age [27,28]. The anticipated region of brain volume coverage by a single sonication, using this specific transducer, was planned based on our in-vivo calibration (Supplementary Figure 1I).

### Fluorescence Imaging

The extracted brains were stored in formalin for at least 24 hours before being evenly sectioned into 10 slices (2 mm slice thickness) for the fluorescence assay. A non-invasive IVIS Spectrum Imaging System (Perkin Elmer, Waltham, MA) was used to record the total amount and distribution of dye molecules delivered to the cisterna magna based on their photon flux. The fluorescence exposure time was set to one second. The F-stop was set at two, and the binning factor was four. The height of the subject was fixed to 0.2 cm. A pair of indocyanine green background, excitation passband: 665-695 nm, and emission passband: 810-875 nm filters were used to filter the fluorescence of IR800 dye molecules. To obtain the IR800 calibration curve, a stock solution was prepared by dissolving 2.0 mg of IR800 in 2.0 mL of DMSO, which was then serially diluted with artificial cerebral spinal fluid to obtain solutions with dye concentrations ranging from 20.0 ug/mL to 0.16 ug/mL as noted in Figure 2D-E. In-vivo calibration of IR800 was also performed upon intrathecal injection of the different sized IR800 tracers (free IR800 i.e., 1kDa, and IR800-labeled IgG antibody i.e., 160 kDa) at different anesthetic states as shown in Figure 2F-I. The fluorescence signal was captured using the IVIS Spectrum Imaging System with the same settings described above.

### Image Processing

After confirming the successful intrathecal injection by observing the R800 signals in whole rat brain IVIS imaging, we proceeded to section the brain into 2 mm slices as shown in Figure 4C and performed additional imaging to quantify the IR800 signals in two distinct manners. Firstly, we measured the average fluorescence intensity, as illustrated in Figure 4A, to determine the amount of IR800 delivery in each brain slice. To calculate this average fluorescence intensity, we computed the mean fluorescence intensity (total flux per second) from the corresponding brain slices of the animals within each specific group. Each data point in our analysis represents the average value of a particular brain slice, ranging from 1 to 10, and encompasses slices from all animals within that particular group. Secondly, we assessed the relative distribution to investigate whether the level of anesthesia impacted the diffusion of molecules within the brain (Figure 4D). Particularly, we aimed to identify which brain slice, corresponding to a specific part of the brain, would be more affected. To perform this calculation, we normalized the fluorescence signals (total flux per second) of each brain slice by the fluorescence signals of the entire brain.

### Statistical Analysis

The physiological and drug distribution data were presented as mean values along with their standard deviations and were visualized using R software. In addition, we employed the non-parametric Mann-Kendall autocorrelation statistical test to investigate whether physiological parameters, including body temperature, heart rate, respiration rate, perfusion, and oxygen level, collected over time exhibited dependencies on their previous time points. This test evaluated the correlation between data points at different time lags, allowing us to detect any underlying systematic trends or patterns in the dataset. The choice of the Mann-Kendall test was deliberate, as it does not rely on assumptions about the data distribution, making it a suitable method for analyzing our time-series physiological data. In assessing these correlations, we used Pearson’s correlation coefficient, which quantifies the strength and direction of relationships between variables. A value of zero indicates no correlation, while positive and negative values indicate positive and negative correlations, respectively. To account for correlations at different time lags, we calculated partial autocorrelations, which remove the effects of shorter-lag correlations. Furthermore, for both physiological and drug distribution data, we conducted statistical analyses employing both one-way and multivariate ANOVA tests. The significance levels were represented using a star rating system: a p-value less than 0.05 was denoted with one star, less than 0.01 with two stars, and p-values less than 0.001 were associated with three stars, demonstrating different degrees of statistical significance.

## Results

To maintain consistent anesthesia levels in the three primary groups (Heavy: anesthesia with 3% isoflurane, Moderate: anesthesia with 2% isoflurane, and Light: anesthesia with 1.5% isoflurane), the flow rate of isoflurane was adjusted according to the animal’s body weight. The body temperature was consistently regulated within the range of 36.5 to 37.5°C throughout the experiments. A thorough monitoring protocol was in place to observe critical physiological indicators in the rats, which included heart rate, respiratory rate, oxygen levels, perfusion, and body temperature. The purpose of this careful observation was to determine if ultrasound had any notable impact on these vital physiological markers, potentially influencing during the glymphatic transport processes. Moreover, experiments were consistently conducted at the same time of day to mitigate potential variations in glymphatic activity due to the rats’ circadian rhythms [29]. For the ultrasound procedure, we followed a protocol based on our prior study. In brief, the protocol involved 0.2 MPa in situ pressure and a local duty cycle of 7.7%, lasting for a total of 10 minutes. This specific ultrasound intensity was chosen because it is significantly low, 10 times below FDA-approved limits for diagnostic ultrasound and can be achieved using existing FDA-approved clinical ultrasound systems, suggesting its ease of translation into clinical applications. It’s noteworthy that the total temperature rise in the sonicated area resulting from this level of ultrasound exposure is estimated to be less than 0.01°C [27]. For sham procedures, the input power to the ultrasound transducer was zero.

### Ultrasonic glymphatic perturbation increases alertness in Light Anesthesia and causes drowsiness in Heavy Anesthesia

To identify any significant patterns or trends in our time series physiological data, we utilized the Mann-Kendall test. As noted in Supplementary Figures 2 and 3, the autocorrelation function (ACF) is close to 1 and partial autocorrelation functions (PACF) are close to zero. A high ACF value close to 1 suggests that each data point is highly dependent on its recent historical values, and this dependence extends to multiple lags. A low PACF value close to zero suggests that the influence of earlier time points has already been accounted for, and at this particular lag, there is little direct relationship between the current value and the value from that specific lag. So, observing a high ACF close to 1 and a low PACF near zero characterizes an autoregressive process. This signifies that a variable heavily relies on its recent history and gradually decreases its dependence on past values as time advances. This understanding helps predict which physiological parameters are impacted solely by ultrasound interventions.

In addition, we examined the average data encompassing key parameters during an approximately 800-second sonication period to understand ultrasound’s influence on vital signs, as depicted in Figure 3. This comparison was made across three anesthesia levels within both Ultrasound and Control conditions using a multivariate ANOVA test. Among these parameters, such as pad temperature regulating body heat within the 36-37°C range, rectal temperature, oxygen levels, heart rate, and perfusion, no statistically significant differences were observed during sonication across various anesthetic sub-groups in both Ultrasound and Control groups, as depicted in Figure 3A. This indicates that these parameters seem unaffected by the tested anesthesia doses and ultrasound application. However, within both the Ultrasound and Control groups shown in Figures 3B and 3D, there was a significant difference in respiratory rate between the 1.5% and 3% anesthesia levels. This discrepancy appears to be influenced by the significant variation in the administered anesthetic doses among these groups. But, uniquely in the Ultrasound group, significant differences were further noted between the 1.5% and 2% anesthesia levels and between the 2% and 3% anesthesia levels (Figure 3D) along with 1.5% and 3% like in Control, suggesting ultrasound’s influence on the respiratory rate pattern.

**Figure 3.**
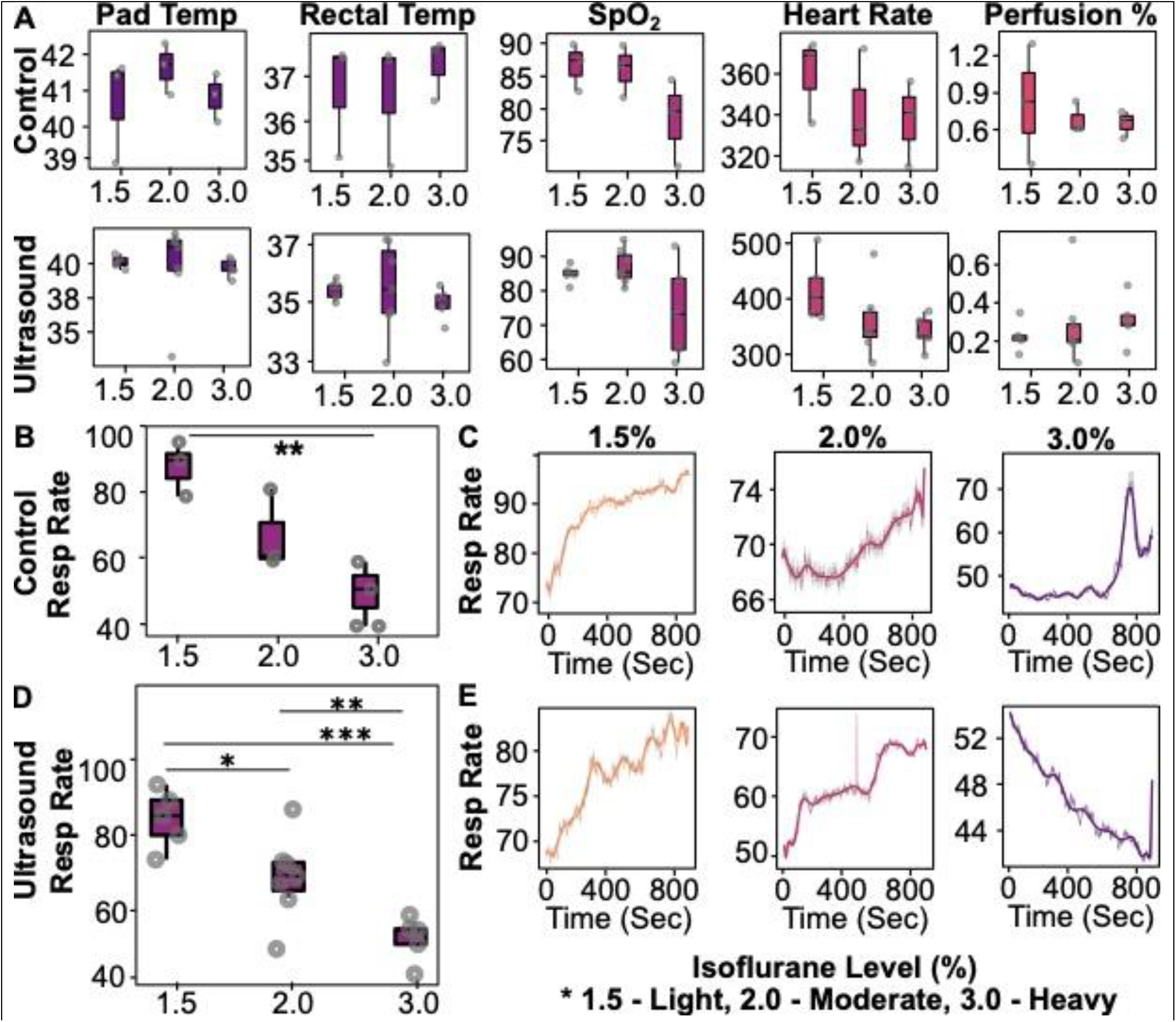
Physiological responses and respiratory patterns under anesthesia with and without ultrasonic influence. **A**. Illustrates average values of various physiological parameters (body temperature, oxygen level, heart rate, perfusion, and pad temperature) measured during sham and ultrasonic sonication at three anesthetic levels: light anesthesia (1.5% isoflurane), moderate anesthesia (2% isoflurane), and heavy anesthesia (3% isoflurane). **B**. Depicts the average respiratory rate at the three different anesthetic conditions in the Control group, accompanied by corresponding respiratory traces in panel **C. D**. Displays the average respiratory rate at the three different anesthetic conditions in the Ultrasound group, with corresponding respiratory traces showcased in panel **E**. In both the Ultrasound and Control groups, significant differences in respiratory rate were observed between 1.5% and 3% anesthesia levels, likely due to varying anesthetic doses. However, the Ultrasound group displayed additional distinctions between 1.5% and 2%, and between 2% and 3%, indicating ultrasound’s influence on respiratory patterns. The Control group consistently showed increased respiratory rate patterns across all anesthesia conditions (**3C**), suggesting anesthesia doesn’t impact this trend. Contrastingly, in the Ultrasound group (**3E**), two distinct respiratory rate patterns emerged. Initially, there was a significant rise in rates, particularly at 1.5% compared to 2%, highlighting ultrasound’s impact on elevating rates during lighter anesthesia. Additionally, a unique trend revealed a decrease in respiratory rate at 3%, significantly different from 2%, suggesting ultrasound might reduce rates as animals move deeper into anesthesia. A multivariate ANOVA t-test was conducted across the groups. Data is represented as mean ± S.D. *: p ≤ 0.05, **: p ≤ 0.01, ***: p ≤ 0.001.

This led to a detailed analysis of respiratory patterns during an 800-second sonication period for both the Ultrasound and Control groups, as illustrated in Figures 3C and 3E. In the Control group (Figure 3C), a consistent rise in respiratory rate occurred across all anesthesia levels. This indicates an unchanging respiratory pattern despite varying anesthesia levels in the Control group. However, in the Ultrasound group, two distinct respiratory rate patterns occurred. Firstly, there was an increase in the respiratory rate at 1.5% and 2% anesthesia levels, notably significant at 1.5% compared to 2%, a contrast absents in the Control group. This highlights ultrasound’s ability to significantly heighten the respiratory rate during lighter anesthesia. Secondly, a decrease in respiratory rate was observed at 3% anesthesia within the Ultrasound group (Figure 3E-Right), a unique trend. A statistically significant difference between 2% and 3% anesthesia levels, particularly evident at 3% compared to 2%, was observed in the Ultrasound group, indicating ultrasound’s role in reducing respiratory rates pattern as animals enter deeper anesthesia. Also, the clear difference between 1.5% and 3% anesthesia levels could be attributed to significant variations in anesthetic doses and the combined impact of ultrasound, potentially contributing to suppressing the respiratory rate. The stronger p-value (p < 0.001) observed in the Ultrasound group compared to the Control group (p < 0.01) when comparing the effects of 1.5% and 3% anesthesia levels suggests that ultrasound intensifies the effects caused by differences in anesthetic doses. Overall, these findings highlight ultrasound’s dual influence: enhancing alertness under lighter anesthesia and inducing drowsiness in heavily anesthetized animals.

Further, it is crucial to emphasize the value of our visual observations in monitoring the recovery process throughout the experiment, with these observations being supported by the physiological data. We noticed that within the Light Anesthesia Ultrasound group, all animals exposed to ultrasound experienced a notably expedited recovery. By the end of the final treatment loop, even as anesthesia was still being administered, they exhibited clear signs of increased alertness. In contrast, in the Light Anesthesia Control group, all animals required a minimum of 3 to 4 minutes after the last mock treatment loop to reach a similar state of wakefulness as the Ultrasound case. A particularly significant finding was that in the Light Anesthesia Ultrasound group, approximately 30% of the animals regained consciousness before completing the final treatment loops, resulting in the early termination of the experiment. In contrast, the animals in the Heavy Anesthesia Ultrasound group experienced a significantly deeper level of anesthesia induced by ultrasound, which prolonged the time needed for them to regain consciousness. This delay amounted to approximately 15-20 minutes longer than the Heavy Anesthesia Control animals, once anesthesia had been discontinued. No noticeable changes were detected in animals under moderate anesthesia levels in both the Control and Ultrasound Groups. Overall, the physiological and observational data suggest that ultrasound prompts distinct effects on animals’ physiological responses, influenced by their initial anesthesia levels. It enhances alertness and expedites recovery in lightly anesthetized animals while it induces drowsiness in heavily anesthetized animals, resulting in a prolonged recovery from ultrasonic glymphatic therapy. However, more investigation is required to gain a complete understanding of how ultrasound influences the recovery process in rats exposed to different levels of anesthesia.

### Efficacy of ultrasound-induced glymphatic transport improves under Light Anesthesia

To evaluate the effects of different anesthesia levels on the efficiency of ultrasonic glymphatic transport of molecules, we conducted experiments with animals under three different isoflurane-induced anesthesia conditions: heavy anesthesia using 3% isoflurane, moderate anesthesia with 2% isoflurane, and light anesthesia with 1.5% isoflurane. In this series of experiments, we chose two imaging tracers with varying molecular sizes: Free IR800-1kDa and IgG – IR800-labeled IgG antibody-160 kDa, to investigate whether the molecule size influenced transport efficiency under these distinct physiological states. We administered intrathecal injections of these imaging tracers. Subsequently, the Ultrasound group received ultrasound exposure within 10 minutes after injection, while the Control group did not receive any ultrasound treatment. We monitored vital signs throughout the experiment. After 70 minutes from the intrathecal injection, we extracted the brains for ex-vivo tissue analysis.

To quantitatively measure the ultrasound-induced movement of IR800 molecules delivered in their free form as IR800 and in their conjugated form as IR800-IgG within the glymphatic system, ex-vivo brain samples were sliced into 2 mm thick sections across the coronal plane of the rat brain, ranging from Bregma-14 to Bregma+4, as illustrated in Figure 4C, and imaged IR800 molecules using IVIS. After conducting ex-vivo imaging of brain slices, we analyzed the average fluorescence intensity for varying anesthesia conditions in both the Ultrasound and Control groups using GraphPad Prism. This comparative analysis across all groups is shown in Figure 4A. To calculate the average fluorescence intensity in Figure 4A, we average the fluorescence intensity (total flux/sec) from the corresponding brain slices of the animals in each specific group. In brief, each data point in our analysis represents the average value of a particular brain slice (ranging from 1 to 10), encompassing slices from all animals in that group. The two-way multivariant ANOVA test shows that the average fluorescence signals were notably higher in the Ultrasound group, particularly in the moderately anesthetized (2%) and heavily anesthetized (3%) for smaller molecule size IR800 groups, compared to their respective Control (no ultrasound) groups. Moreover, animals under moderate anesthesia exhibited a significantly lower p-value (p = 0.0014) when contrasted with heavily anesthetized counterparts (p = 0.0233), indicating that ultrasound is more effective in transporting smaller size molecules toward the lower anesthetic domain (Moderate group) compared to a state of higher anesthesia (Heavy group).

**Figure 4.**
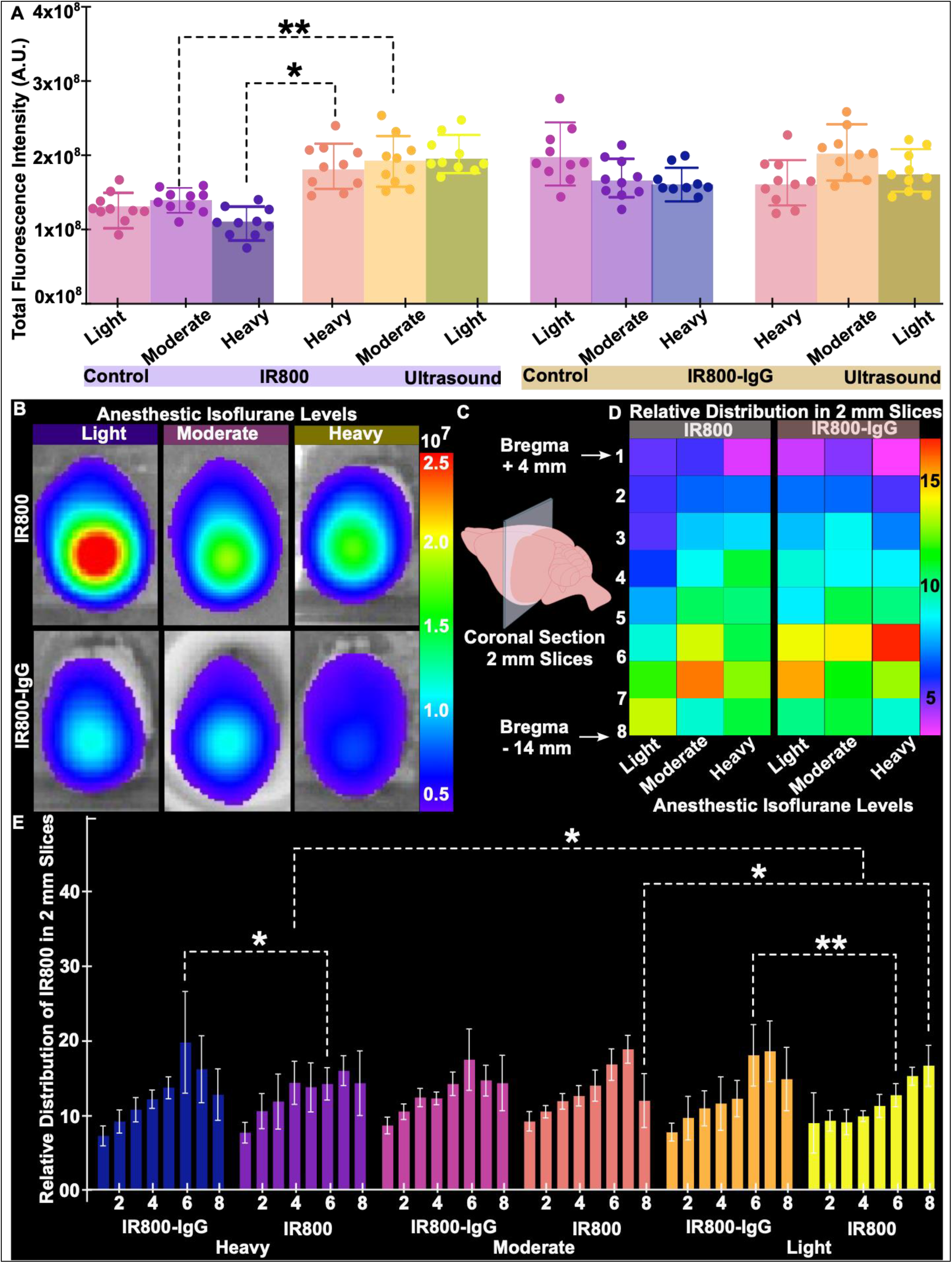
Ultrasound’s Impact on Glymphatic Transport Efficiency. **A**. Comparative Analysis: Shows higher fluorescence signals in the Ultrasound group, especially in moderately and heavily anesthetized states for smaller IR800 molecules. **B**. Ex-vivo IVIS Image: Illustrates rat brain exposure to various-sized IR800 agents and isoflurane anesthesia levels in Ultrasound group. **C**. Brain Slicing for IVIS: Illustration of rat brain sections for IR800 quantification. **D**. IR800 Distribution Visualization: Depicts IR800 signals across the rat brain subjected to ultrasound, using a heatmap spanning from Bregma-14 to Bregma+4. **E**. Statistical Analysis: Highlights faster transport of smaller IR800 molecules, particularly in lighter anesthesia states. Indicates efficient ultrasonic transport via the glymphatic system, especially with smaller molecules. A multivariate ANOVA t-test was conducted across the groups. Data is represented as mean ± S.D. *: p ≤ 0.05, **: p ≤ 0.01.

However, in the Light anesthetized Ultrasound group, the average fluorescence intensity showed no statistically significant difference compared to the Control group, irrespective of molecule size, presenting challenges in interpreting the data. The lack of statistical significance could be due to several factors. It is possible that under Light anesthesia, ultrasound’s robust influence on molecule transport swiftly moves substances from the injection site (cisternal magna) as compared to Moderate and Heavy cases. This rapid motion might efficiently cleanse the brain, allowing molecules to exit the glymphatic system and enter the peripheral lymphatic system. This emphasizes the importance of focusing on the detailed analysis of molecules’ presence in individual slices, starting from the tracer injected site all the way to the olfactory bulb, a major part of existing tracers from glymphatic to the lymphatic system, and performing slice-by-slice comparisons between different anesthetic levels in Ultrasound group. So, we performed relative distribution. The relative distribution of IR800 molecules involved normalizing the fluorescence signals of each brain slice by the fluorescence signals of the entire brain obtained from the whole brain IVIS image. Figure 4B shows a representative ex-vivo IVIS image of rat brains after exposure to the same level of ultrasound but different-sized IR800 agents intrathecally, along with various levels of isoflurane anesthesia. Figure 4B was used to calculate the total fluorescence intensity. These brain samples were subsequently sliced into 2 mm thick sections spanning from Bregma-14 to Bregma+4 across the coronal plane of the adult rat brain, as shown in Figure 4C. Figure 4D showcases a heatmap illustrating the normalized distribution of IR800 within 2 mm-thick brain slices, while 4E shows a slice-by-slice comparison of IR800 distribution across different anesthetic levels and molecular sizes.

The heat map demonstrates that within the free IR800 molecules, the Light Anesthesia group consistently exhibits a stable fluorescence signal intensity of IR800 molecules, ranging from a maximum of light-yellow level and a minimum of light purple. Conversely, all other groups showcase signals that either exceed the maximum, indicating tracer aggregation instead of diffusion towards adjacent slices, or fall below the minimum, implying insufficient tracer presence, in at least one slice per group. This suggests that when the brain is under light anesthesia, ultrasound effectively increases the movement of molecules within the brain-fluid space. This increased facilitation boosts the chances of these molecules reaching the exit points from the glymphatic to lymphatic sites, which ultimately speeds up their clearance. Consequently, this could potentially decrease the overall presence of IR800 molecules throughout the entire brain system. Further statistical analysis supports that notion in Figure 4E. In the Light Anesthetic group, within the 6th slice closer to the injected space (cisternal magna), the smaller molecular size of IR800 presence a statistically significantly lower as compared to the larger size IR800. This pattern is evident in the Heavy Anesthesia Group as well, where the smaller IR800 molecule accumulates significantly less in the 6th slice compared to the larger one. This indicates that smaller molecules are transported more swiftly away from the cisterna magna than larger ones. Additionally, within the smaller IR800 molecule, the 5th slice in the Lighter Anesthetic Group exhibits notably lower accumulation compared to the Heavy Group. This suggests that smaller molecules are transported faster from the cisterna magna in a Lighter Anesthetic state compared to a Heavy one. Overall, lighter anesthesia and smaller-sized tracers might demonstrate improved transport efficiency following ultrasonic glymphatic intervention. However, adjusting ultrasound parameters further, possibly by exploring frequencies other than the 650 kHz used in this study, could offer important insights into understanding ultrasound’s limitations concerning molecule size dependency and the influence of different anesthetic levels on the effectiveness of glymphatic transport and clearance.

## Discussion

This study highlights the correlation between the level of anesthesia and the impact of ultrasound-induced glymphatic manipulation on alertness, recovery, and molecule transport. In animals with light anesthesia, it enhances alertness and accelerates recovery resulting in improved molecule transport. Conversely, heavily anesthetized subjects experience increased drowsiness and extend the time required for animals to regain consciousness, leading to a reduction in molecule transport efficiency. The physiological data focuses on the distinct impact of ultrasound therapy on animals’ physiological responses, depending on their initial anesthesia levels. The statistically significant differences in respiratory rate between Light and Heavy anesthesia levels, which were observed both in the Ultrasound and Control groups (Figures 3B & 3D), can be attributed to the substantial variation in isoflurane levels. However, in the Ultrasound group, the significant difference in respiratory rate between Light and Moderate anesthesia levels, as well as between Moderate and Heavy anesthesia levels, suggests that ultrasound has an impact on that (Figure 3D). Further, the recovery process during the experiment was closely monitored and supported by the physiological data. In the Light Anesthesia Ultrasound group, animals exposed to ultrasound exhibited a notably expedited recovery and increased alertness, with some animals regaining consciousness even before the final treatment loops were completed. In contrast, the Light Anesthesia Control group animals required several minutes to reach a similar state of wakefulness. The Heavy Anesthesia Ultrasound group experienced a delay in regaining consciousness compared to the Heavy Anesthesia Control group, indicating that ultrasound-induced anesthesia is more prolonged. Both, physiological and recovery data suggest that ultrasound heightens the alertness in animals under light anesthesia, while it promotes drowsiness in heavily anesthetized animals. Several studies have highlighted the significance of respiration, aquaporin 4 (AQP4) expression, body posture, and vasomotor activities in influencing the functionality of the glymphatic system, as it naturally varies based on these states [30]. Our physiological data additionally indicate that ultrasound can further modulate glymphatic functionality by influencing respiratory patterns. This finding demands for careful consideration when utilizing ultrasound-related brain applications, particularly in ongoing preclinical and clinical settings. Moreover, this study underscores ultrasound’s potency as a tool to investigate brain functionality, emphasizing the extensive time and effort required for a comprehensive understanding of its effects.

Moreover, our data on ultrasound’s influence on drug movement through the glymphatic system, particularly in different physiological states, demonstrates that employing lighter to moderate anesthesia is more beneficial than heavy anesthesia, particularly for molecules approximately 1kDa in size (as depicted in Figure 4A). The rationale behind this enhanced movement of drug molecules lies in ultrasound’s impact on altering animals’ breathing patterns, increasing it under lighter to moderate anesthesia and decreasing it under heavy anesthesia. These changes in breathing patterns influence fluid flow around the brain, allowing varying amounts of injected IR800 molecules to penetrate brain tissue. This highlights the significant impact of altered breathing patterns induced by ultrasound on fluid movement in the brain. This concept is supported by a study using mathematical modeling, indicating that even slight changes in pressure due to breathing have a larger effect on brain fluid flow than other physiological parameters like the heart’s pumping action [31]. Incorporating this study strengthens the idea that respiratory pattern plays a vital role in cerebrospinal fluid movement and warrants further investigation into other underlying mechanisms in cellular-molecular. One avenue worth exploring involves investigating how ultrasound affects the expression of aquaporin 4 (AQP4) at astrocytic endfeet, responsible for fluid exchange. Another intriguing area of study is examining the brain’s respiratory electron transport chain.

Yet, our interpretation of some drug transport data encounters limitations. In the Light Anesthesia Ultrasound group (Figure 4A), the average fluorescence intensity displayed no statistically significant difference in comparison to the control group, regardless of molecule size which poses challenges in interpretation. The absence of statistical significance in this context could potentially be attributed to a couple of factors. It’s possible that under conditions of Light Anesthesia, molecules influenced by ultrasound experience faster clearance in comparison to Moderate and Heavy anesthesia levels. Alternatively, it could be that in some animals, the experiment had to be terminated prematurely due to the animals regaining consciousness more quickly as a result of the ultrasound therapy, which may not have provided adequate exposure. However, a more detailed examination of clearance kinetics could provide further insights, though it falls outside the scope of this study. Additional factors that may contribute to the absence of statistically significant results could include a relatively small sample size or variations in experimental variables such as intrathecal injection, animal size, tracer handling, imaging procedures, and other potential confounding factors. Another study revealed that incorporating pain medication during isoflurane-induced anesthesia notably boosted the fluid around brain cells, leading to quicker glymphatic system transport [32]. This implies a potential synergistic effect in the isoflurane-induced anesthetic state when combined with neuromodulatory molecules. This prompts us to question whether ultrasound, with its mechanical neuromodulation capability [33–36], can have a synergistic effect with isoflurane-induced anesthesia. It’s important to note that our glymphatic manipulation ultrasound pressure and sonication time differ from the neuromodulation effects reported in preclinical and clinical studies. However, these specific aspects have not been explored in prior ultrasound applications, which introduces a new avenue for ultrasound research.

## Conclusions

Overall, our study shows that ultrasound’s impact is linked respiratory pattern depend on the level of anesthesia: it enhances alertness in lightly anesthetized animals during ultrasonic glymphatic therapy, which improves glymphatic transport, while it induces drowsiness in heavily anesthetized animals and a subsequent reduction in glymphatic transport. The results suggest that a low level of anesthesia is beneficial for efficient ultrasonic glymphatic transport, making it more in line with the awake state in current clinical ultrasound treatments, thus advancing the technology closer to clinical translation. However, a more in-depth examination and analysis are necessary to fully uncover the underlying mechanisms contributing to these observed results.

## Supporting information

AryalSuppleMethod

## Acknowledgment

We extend our gratitude to the In Vivo Imaging Centre at Stritch School of Medicine at Loyola University Chicago for their generous provision of the IVIS system and access to the animal facility. We also appreciate the contributions and support of all members of the Aryal Lab, whose insightful discussions and equipment usage were invaluable. Gabriel Gallegos, Nuala Kalensky, Dhruv Patel, and Esther Wayntraub are greatly appreciated for their work in designing the figure’s schematics. Special thanks are due to Dr. Maurizio Bocchetta and Dr. Kelly Langert for their valuable insights into optical imaging. Financial support for this work was provided by the Engineering Department’s Faculty Start-Up fund at Loyola University Chicago, the Department of Chemical, Biological, and Bioengineering at North Carolina Agricultural and Technical State University, as well as support from the National Academy of Medicine Healthy Longevity Catalyst Award and the Focused Ultrasound Foundation Global Intern program.

