## Supplementary material for "The Impact of Anesthesia on Ultrasonic Glymphatic Perturbation Protocol: Alertness, and Glymphatic Influx": AryalSuppleMethod

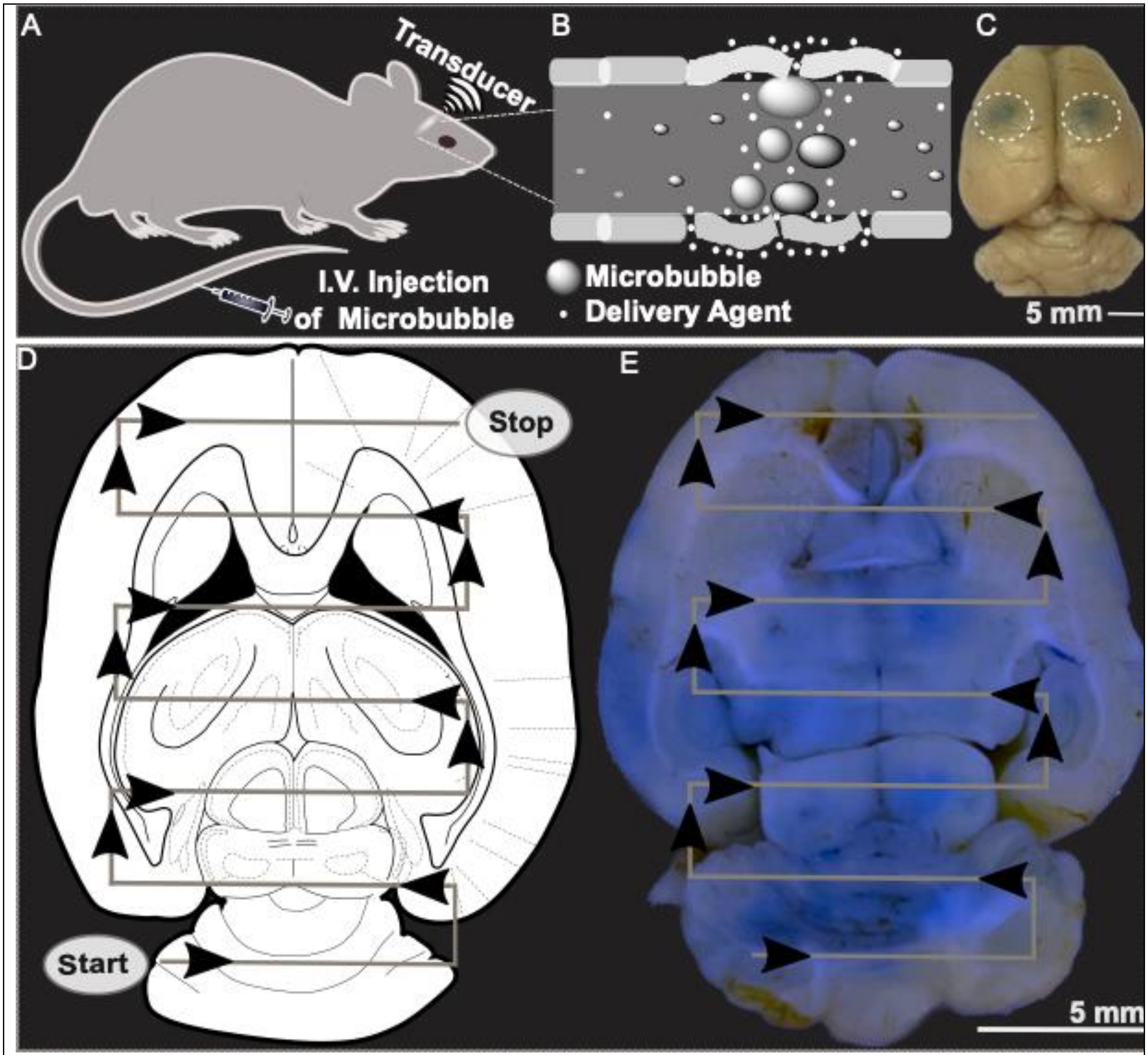


Supplementary Figure 1. *Preclinical transcranial focused ultrasound system calibration in in-vivo setting using blood-brain barrier-based drug delivery assay. To confirm the ultrasound field's behavior in in-vivo glymphatic study, we implemented the vascular barrier opening assay using an established ultrasound pulse (650 kHz, 1PRF, 1 min, 0.35 MPa), in combination with an ultrasound contrast agent, evaluated trypan blue diffusion in ex-vivo rat brain slice (E).*

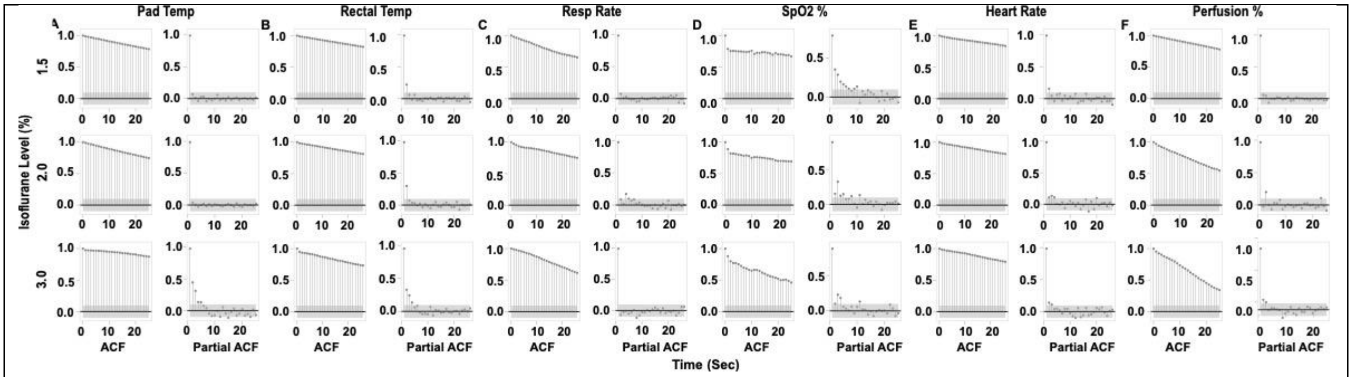


***Supplementary Figure 2.*** *Control (no ultrasonic glymphatic manipulation). The autocorrelation function (ACF) is close to 1 and partial autocorrelation functions (PACF) are close to zero. Non-parametric Mann-Kendall autocorrelation statistical test to investigate whether physiological parameters, including body temperature, heart rate, respiration rate, perfusion, and oxygen level, collected over time exhibited dependencies on their previous time points.*

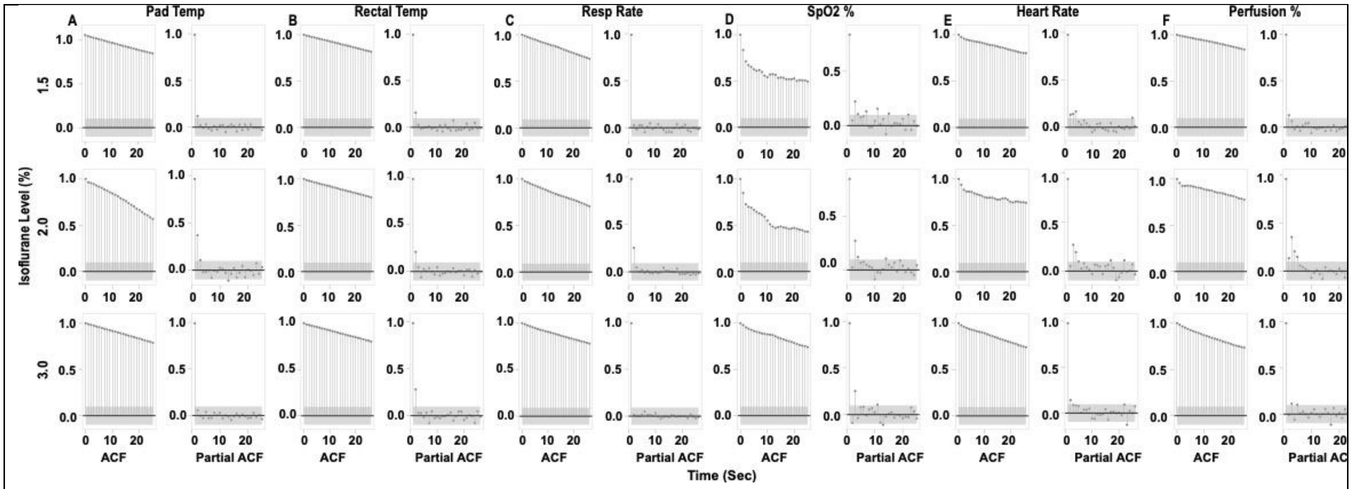


**Supplementary Figure 3.** *Ultrasonic glymphatic manipulation Groups: The autocorrelation function (ACF) is close to 1 and partial autocorrelation functions (PACF) are close to zero. Non-parametric Mann-Kendall autocorrelation statistical test to investigate whether physiological parameters, including body temperature, heart rate, respiration rate, perfusion, and oxygen level, collected over time exhibited dependencies on their previous time points.*
